# GABA mediates experience-dependent regulation of myelination in the mouse visual pathway

**DOI:** 10.1101/2025.03.05.641594

**Authors:** Yasuyuki Osanai, Batpurev Battulga, Reiji Yamazaki, Kenta Kobayashi, Kenji Kobayashi, Yuka Nakamura, Masaki Ueno, Hiroaki Mizukami, Yumiko Yoshimura, Nobuhiko Ohno

## Abstract

Myelination in the visual pathway is critical for transmitting visual information from retina to the brain. Reducing visual experience shortens myelin sheath length and slows the conduction velocity of the optic nerve. However, the mechanism underlying such experience-dependent myelination is unclear. Here, we found that closing both eyes, binocular deprivation (BD), during the juvenile period less affects the optic nerve myelination than monocular deprivation (MD) via GABA signaling. RNA-seq analysis of optic nerves from MD and BD mice revealed that GABAergic signaling is downregulated on the deprived side of MD compared to the intact side and BD. Inhibition of GABAergic signaling during the juvenile period resulted in myelin sheath shortening and excessive oligodendrocyte generation in normal mice, similar to the changes observed in MD mice. Enhancing GABAergic signaling rescued the myelin sheath shortening and excessive oligodendrocyte generation in the optic nerve of MD mice. Furthermore, we identified novel GABAergic neurons located within the optic nerve, whose neurites form belt-like presynaptic structures with the oligodendrocyte lineage cells, suggesting a potential source of the GABAergic inputs into oligodendrocytes. Our results indicate that the myelination of visual pathway is maintained by binocular visual inputs via intra-nerve GABA signaling.

## Introduction

Myelination facilitates nerve conduction and is essential for brain functions. The structural changes of myelin sheaths depending on sensory experiences can modulate the nerve conduction velocity and affect the functions of the neural circuitry^1,2^. Optic nerve axons are fully myelinated^3,4^, and their myelination is necessary for accurate visual perception^5^. Visual symptoms are often the earliest manifestation of demyelinating diseases such as multiple sclerosis^6,7^. The myelination of the optic nerve is regulated by both neuronal activity-dependent and cell-autonomous manner. Seminal work of Raff and Barres revealed that proliferation of oligodendrocyte progenitor cells (OPCs) in the rat optic nerve is dependent on a growth factor secreted in response to retinal neuronal activity, established the concept of activity dependent myelination^8^. Subsequently, proliferation of OPCs depending on neuronal activity has been well characterized in multiple methods^9,10^. However, differentiation from OPCs to mature oligodendrocytes does not appear to require neuronal activity in the optic nerve, as shown by rodent models where the eye was removed^11,12^. Instead, visual input has been shown to play a crucial role in inducing morphological changes in oligodendrocytes and myelin in the visual pathway. Studies have revealed that monocular deprivation (MD), closing one eye through eyelid suturing, reduces myelin sheath length in the optic nerve and optic chiasm and slows nerve conduction velocity^13,14^. In addition, MD affects remodeling of myelination in the visual cortex, especially for Parvalbumin positive GABAergic neurons^15^. Thus, it is possible that the low visual acuity of deprived eye in MD can be partly attributable to the impairment of myelination.

MD shifts ocular dominance toward the non-deprived eye in the visual cortex^16,17^ and alters axonal structure in dorsal lateral geniculate nucleus^18,19^. In contrast, despite losing visual input from both eyes, mice that experience binocular deprivation (BD) during the visual critical period exhibit better visual acuity than those undergo MD and show ocular dominance similar to normally reared mice^20,21^. Indeed, one treatment for anisometropic amblyopia, a condition characterized by low visual acuity in one eye, is patching the health eye^22^. However, if reduction of visual input impairs normal myelination as observed in MD, myelination in BD will be perturbed significantly. Conversely, the improved visual function in BD argues that the changes in myelination and nerve conduction in visual pathway may be ameliorated by BD. However, little is known about BD effect on myelination of visual pathway, and the mechanistic background in myelination dependent on visual input is unclear.

In this study, we leveraged whole mount optic nerve/optic chiasm immunohistochemistry with genetic oligodendrocyte labeling and found that BD does not affect myelination of visual pathway. Based on the apparent discrepancy in the effects of visual input on myelination in MD and BD, differential gene expression analysis using RNA-seq was performed, revealing that GABAergic signaling is downregulated in the optic nerve of deprived eye in MD but not in BD. Consistently, inhibition of GABAergic signaling recapitulated the changes in oligodendrocyte number and myelin morphology observed in deprived side of MD, while enhancement of GABAergic signaling rescued the changes of oligodendrocytes and myelin observed in MD mice. Furthermore, we identified novel GABAergic neurons in the optic nerve as a potential source of GABA. Our results indicate that myelination of optic nerve is regulated by the balance of visual inputs from both eyes, similar to neuronal plasticity in the visual cortex, via GABAergic signaling.

## Result

### Myelination of the visual pathway remains intact in BD mice

Longer myelin sheaths increase nerve conduction velocity in the optic nerve^13,23^. Oligodendrocytes in the optic chiasm surrounded by axons from input-deprived eye (hereafter MD deprived oligodendrocytes) form shorter myelin sheaths compared to those surrounded by axons from intact eye (hereafter MD intact oligodendrocytes) in the MD mice^24^. To investigate whether oligodendrocytes in the BD form short myelin sheaths, we visualized oligodendrocytes in the BD mouse optic chiasm using rabies virus as reported previously^25^, and compared their morphology to those in MD mice observed in the report^24^ (Figure 1**a**). Interestingly, the length of myelin sheath formed by oligodendrocytes in BD mice was significantly longer than that of MD deprived oligodendrocytes, and similar to those formed by MD intact oligodendrocytes and oligodendrocytes equally surrounded by intact and deprived axons in MD mice (hereafter MD intermingled oligodendrocytes) (Figure 1**b** – 1**d**). The number of processes formed by individual oligodendrocytes tended to be low in MD intact and BD oligodendrocytes compared to MD deprived and MD intermingled oligodendrocytes (Figure 1**e**). If individual oligodendrocytes form significantly shorter myelin sheaths, an excessive number of oligodendrocytes would be required to fully myelinate visual pathway axons (Figure S1**a**). Indeed, the density of CC1^+^Olig2^+^ mature oligodendrocytes tended to be higher in MD deprived optic nerve compared to MD intact optic nerve (Figure S1**b**), consistent with the previous finding^13^. By contrast, the mean number of mature oligodendrocytes as well as oligodendrocyte progenitor cells (CC1^-^Olig2^+^), astrocytes (S100B^+^PDGFRa^-^) and microglia (Iba1^+^) in BD optic nerve was similar to that of MD intact optic nerve (Figure S1**c**-S1**h**). These data indicate that the morphology including myelin sheath and the number of BD oligodendrocytes are similar to those of MD intact oligodendrocytes.

**Figure 1.**
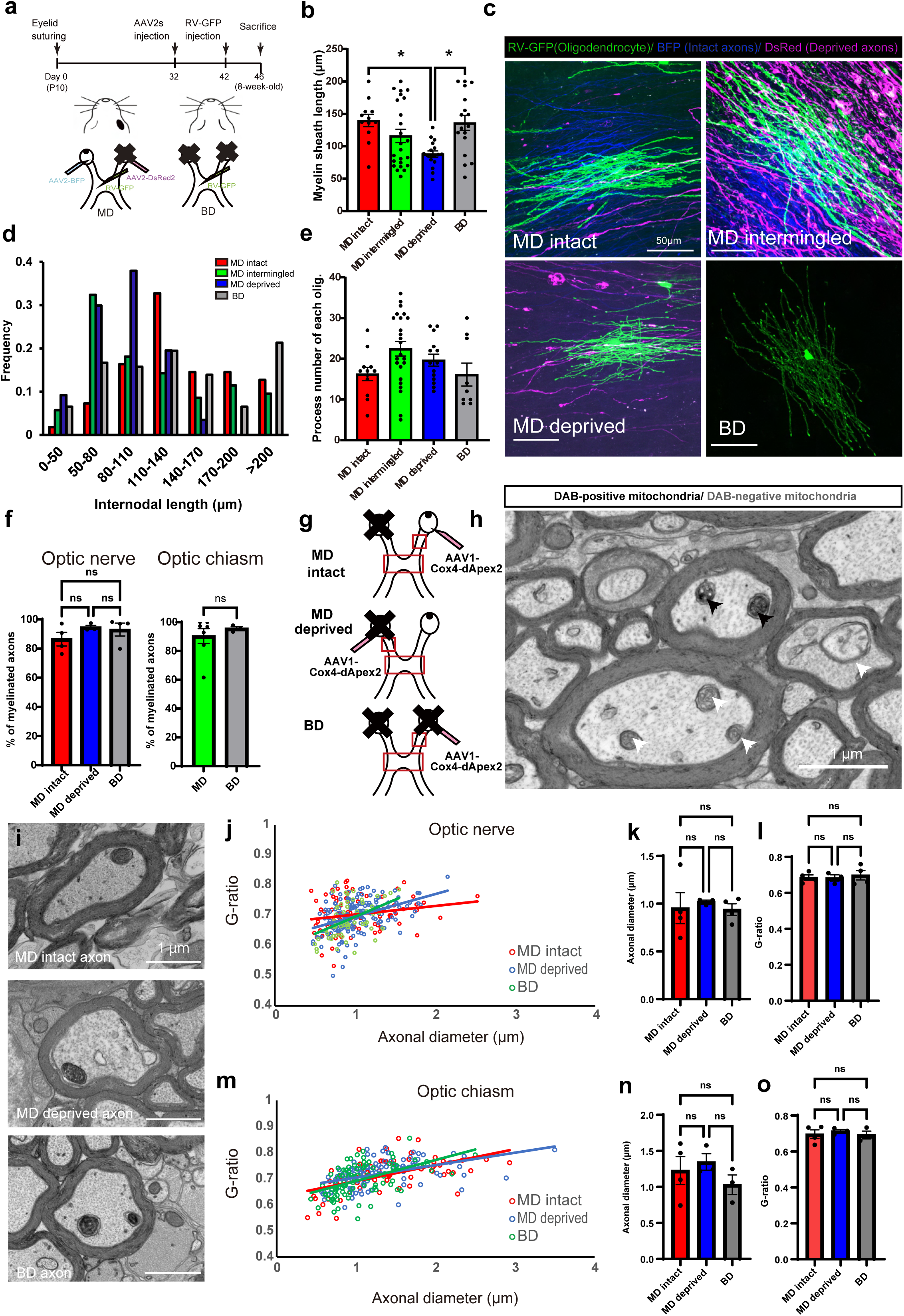
Myelination of BD mice is normal. **a**, Scheme of MD, BD, and viral injection. **b**, Comparison of the average internodal length produced by chiasmal oligodendrocytes in the MD intact, intermingled, deprived region, and BD (MD intact 139.8 ± 9.5 µm, MD intermingled 116.1 ± 10.2 µm, MD deprived 88.0 ± 4.8 µm, BD 136.0 ± 11.5 µm, p = 0.0048). Dots represent mean myelin sheath length of individual oligodendrocytes. **c**, Representative Z-stacked confocal images of oligodendrocytes in the intact (upper left), intermingled (upper right), deprived (bottom left) regions of the MD mouse optic chiasm, and in BD mice (bottom right). **d**, The histogram showing the distribution of myelin sheath length (n = 55, 105, 87, and 108 myelin sheaths, for MD intact, intermingled, deprived, and BD, respectively). **e**, Comparison of the average number of myelin sheaths produced by the individual oligodendrocytes (MD intact 16.3 ± 5.6, MD intermingled 22.5 ± 1.8, MD deprived 19.6 ± 1.5, BD oligodendrocyte 16.1 ± 2.8, p = 0.0465). **f**, Electron microscopic analysis of percentages of myelinated axons. Left: percentage of myelinated axons in the MD intact, deprived, and BD optic nerves (MD intact 86.4 ± 4.6 %, MD deprived 94.5 ± 1.4 %, BD 93.0 ± 4.4 %, p = 0.38). Right: percentage of myelinated axons in the MD and BD optic chiasm (MD 90.4 ± 5.23 %, BD 95.6 ± 1.32 %, p = 0.54). Dots represent the values of individual optic nerves or optic chiasms. **g**, Scheme of MD, BD, and viral injection. **h**, A representative electron microscopic image showing myelinated axons with DAB-positive (black arrowhead) and -negative (white arrowhead) mitochondria in the optic chiasm. **i**, Representative electron microscopic images showing myelinated axons with DAB-positive mitochondria in MD intact axons, MD deprived axons, and BD axons in the optic chiasm. **j**, A scatter plot of G-ratio and axonal diameter in the optic nerve. Dots represent the values of individual dApex2-positive axons. **k, l**, Comparison of the average axonal diameter (**k**) and G-ratio (**l**) from MD intact, deprived, and BD optic nerve (axonal diameter; MD intact, 0.95 ± 0.16 µm; MD deprived, 1.02 ± 0.02 µm; BD, 0.94 ± 0.06 µm; n = 4, 3, 4, respectively, p = 0.88: G-ratio; MD intact, 0.686 ± 0.015; MD deprived, 0.683 ± 0.0178; BD, 0.698 ± 0.0262; n = 4, 3, 4, respectively, p = 0.86). **m**, A scatter plot of G-ratio and axonal diameter in the optic chiasm. Dots represent the values of individual dApex2-positive axons. **n**,**o**, Comparison of the average axonal diameter (**n**) and G-ratio (**o**) from MD intact, deprived, and BD optic nerve (axonal diameter; MD intact, 1.28 ± 0.19 µm; MD deprived, 1.35 ± 0.12 µm; BD, 1.03 ± 0.14 µm; n = 4, 3, 3, respectively, p = 0.48: G-ratio; MD intact, 0.696 ± 0.024; MD deprived, 0.713 ± 0.010; BD, 0.692 ± 0.021; n = 4, 3, 3, respectively, p = 0.78). Data were analyzed by Kruskal-Wallis test with post hoc Dunn’s test (**b**, **e**), by One-way ANOVA with Tukey post hoc test (**f** left, **k**, **l**, **n**, **o**), or by Mann-Whitney U test (**f** right). ns *P* > 0.05; \**P* < 0.05. Bar graphs show mean values, and error bars indicate SEM.

Next, we investigated whether MD or BD influences percentage of myelinated axons and myelin thickness using transmission electron microscopy. Most of the axons in MD and BD mice were myelinated both in the optic nerve and the optic chiasm (Figure 1**f**). To analyze G-ratio and diameter of the axons derived from intact and deprived eyes, AAV1-Cox4-dApex2 was injected into intact or deprived eyes in the MD mice, as well as the eyes in BD mice (Figure 1**g**). Cox4-dApex2 labels mitochondria in axons of AAV-injected eyes (Figure 1**h** and 1**i**)^26^, allowing the identification and analysis of labeled axons from the intact and deprived eyes. This analysis revealed that G-ratio and diameter of BD axons were similar to that of MD intact and deprived axons both in the optic nerve and the optic chiasm, while the axon diameter in the BD tended to be smaller without statistical significance (Figure 1**j** – 1**o**). These data support the concept that myelination of BD is intact.

Mice subjected to BD in juvenile period are known to have better visual functionality compared to those undergo MD^21^. To address if the visual function of BD mice with intact myelination is preserved, we performed the visual cliff test^27,28^, which measures the time mice spend on the shallow area compared to the seemingly deep area (Figure S2**a** and S2**b**). We found that control and BD mice stayed longer on the shallow floor, whereas some MD mice did not (Figure S2**c**). The locomotor activity of the mice was similar among the three groups (Figure S2**d**). Although BD mice did not immediately return to the shallow side upon reaching the border, BD mice stayed longer time in shallow area than deep area (Figure S2**e** and S2**f**). These results indicate that the visual function is well preserved in BD mice and consistent with its intact myelination in the visual pathway.

### Increasing neuronal activity of retinal ganglion cells lengthens myelin sheath in the optic nerve of BD mouse

Since myelination remains intact in BD but is perturbed in MD deprived side, it is possible that unilaterally increasing the neuronal activity of retinal ganglion cells (RGCs) in BD mice shortens myelin sheath in less active axons or elongates it in highly active axons. To address these hypotheses, the neuronal activity of RGC in one eye of BD mice was increased by the injection of AAV2-hM3Dq-mCherry (Figure 2**a**). We confirmed that immunoreactivity of c-Fos, immediate early protein, in RGC of eyes injected with AAV2-hM3Dq-mCherry was higher than in eyes injected with AAV2-BFP, two hours after clozapine-N-oxide (CNO) administration (Figure 2**b**–2**d**). Under the unilateral intraocular injection of AAV2-hM3Dq-mCherry, the axons from either side eyes were sparsely labeled with AAV2-GFP, and the length of myelin sheaths was determined by measuring the distance between the nodes of Ranvier (marked by Caspr immunostaining pairs) on the GFP-labeled axons in the cleared tissue (Figure 2**e**–2**g**). Almost all the GFP-labeled RGC axons from the eyes co-injected with AAV2-GFP and AAV2-hM3Dq-mCherry were mCherry-positive (Figure 2**h**). In the optic nerve, the myelin sheath length of axons from the hM3Dq-mCherry expressing eyes was significantly longer than that of axons from eyes without hM3Dq-mCherry expression, indicating enhancing neuronal activity increases internodal length (Figure 2**i**). In the optic chiasm, where axons derived from AAV2-hM3Dq-mCherry injected eye (ipsilateral) and those from non-injected eye (contralateral) are intermingled, myelin sheath length was similar between those axons (Figure 2**j**), suggesting that myelin sheath length is determined by neuronal activity of both eyes in the optic chiasm. Collectively, increasing neuronal activity in one eye in BD lengthens the myelin sheath of optic nerve. However, this does not shorten the myelin sheath of less active axons, failing to replicate the shorter myelin sheath observed in MD.

**Figure 2.**
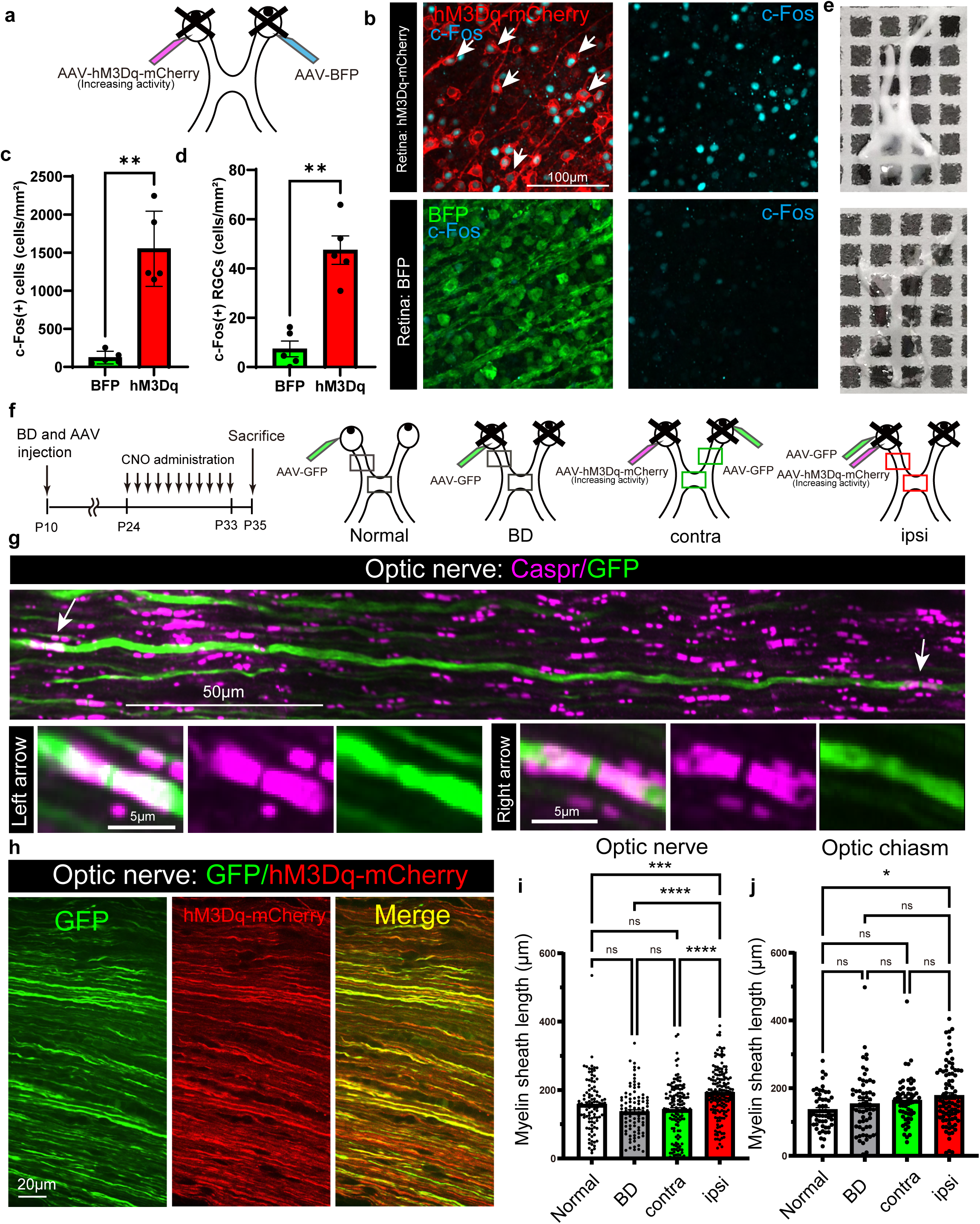
Enhancing neuronal activity increases myelin sheath length. **a**, Scheme of AAV2-hM3Dq-mCherry and AAV2-BFP injection into the eyes of BD mice. **b**, Confocal images of c-Fos-expressing cells in the retina transduced with AAV2-hM3Dq-mCherry or AAV2-BFP. **c**,**d**, Density of c-Fos-positive cells or c-Fos-positive cells with RGC morphology in retina transduced with AAV2-hM3Dq-mCherry or AAV2-BFP (c-Fos-positive cells; BFP, 122.6 ± 37.1 cells/mm^2^; hM3Dq, 1551.0 ± 220.4 cells/mm^2^; p = 0.003: c-Fos-positive RGC; BFP, 7.4 ± 3.2 cells/mm^2^; hM3Dq, 47.5 ± 5.8 cells/mm^2^; p = 0.004). **e**, Representative images of the optic nerve and optic chiasm before (top) or after (bottom) tissue clearing. The grid represents 1 mm square. **f**, Scheme of the experimental schedule and experimental groups. AAV2-GFP was injected into the eye of normal mice, BD mice, or AAV2-hM3Dq-mCherry injected eye (ipsilateral) or non-injected eye (contralateral) of BD mice. **g**, A stitched z-series confocal image showing Caspr signals (magenta) on GFP-expressing RGC axons (green). Arrows indicate the positions of nodes of Ranvier, flanked by the Caspr signals. **h**, Representative images of RGC axons in the optic nerve from a retina co-injected with AAV-GFP and AAV-hM3Dq-mCherry. **i**,**j**, Myelin sheath length of axons in normally reared mice (without CNO injection), BD + CNO mice, BD + hM3Dq + CNO mice contralateral or ipsilateral side in the optic nerve (**i**) and optic chiasm (**j**) (optic nerve; normal 157.2 ± 7.7 µm, BD 135.3 ± 6.7 µm, BD-hM3Dq contralateral 141.7 ± 6.6 µm, BD-hM3Dq ipsilateral 191.7 ± 5.6 µm, p < 0.0001: optic chiasm; normal 135.2 ± 7.5 µm, BD 152.3 ± 11.1 µm, BD-hM3Dq contralateral 162.7 ± 8.3 µm, BD-hM3Dq ipsilateral 176.9 ± 9.7 µm, p = 0.0266). Dots represent the values of individual myelin sheaths. Data were analyzed by paired t-test (**c**, **d**) or by Kruskal-Wallis test with post hoc Dunn’s test (**i**, **j**). ns *P* > 0.05; \**P* < 0.05; \*\**P* < 0.01; \*\*\*\**P* < 0.001; \*\*\*\**P* < 0.0001. Bar graphs show mean values, and error bars indicate SEM.

### GABAergic signaling regulates oligodendrocyte differentiation and its morphology

To find the mechanisms of shorter myelin sheaths in MD deprived side and normal myelin sheaths in BD, we performed bulk RNA-seq with optic nerves obtained from MD deprived side, MD intact side, and BD (Figure 3**a**). As there are no RGC cell bodies in the optic nerve, difference of gene expression detected by RNA-seq will be derived from cells in the optic nerve such as oligodendrocyte lineage cells, astrocytes, and microglia. We measured 24346 genes, and detected 52 genes that were downregulated (fold changes > 2.0) in MD deprived side compared to MD intact side. Of those, 29 genes were also down regulated in MD deprived side compared to BD (fold changes > 1.2) (Figure 3**b**). Unsupervised analysis of Gene Ontology for these genes using Metascape^29^ revealed that genes related to GABA receptor activation, memory, and cell cycle check points related genes were significantly down regulated in MD deprived side compared to MD intact and BD (Figure 3**c**). We further searched the Barres Brain RNA-seq database^30,31^ and found that these genes related to GABAergic receptor activation are expressed in oligodendrocyte lineage cells (Figure 3**d**).

**Figure 3.**
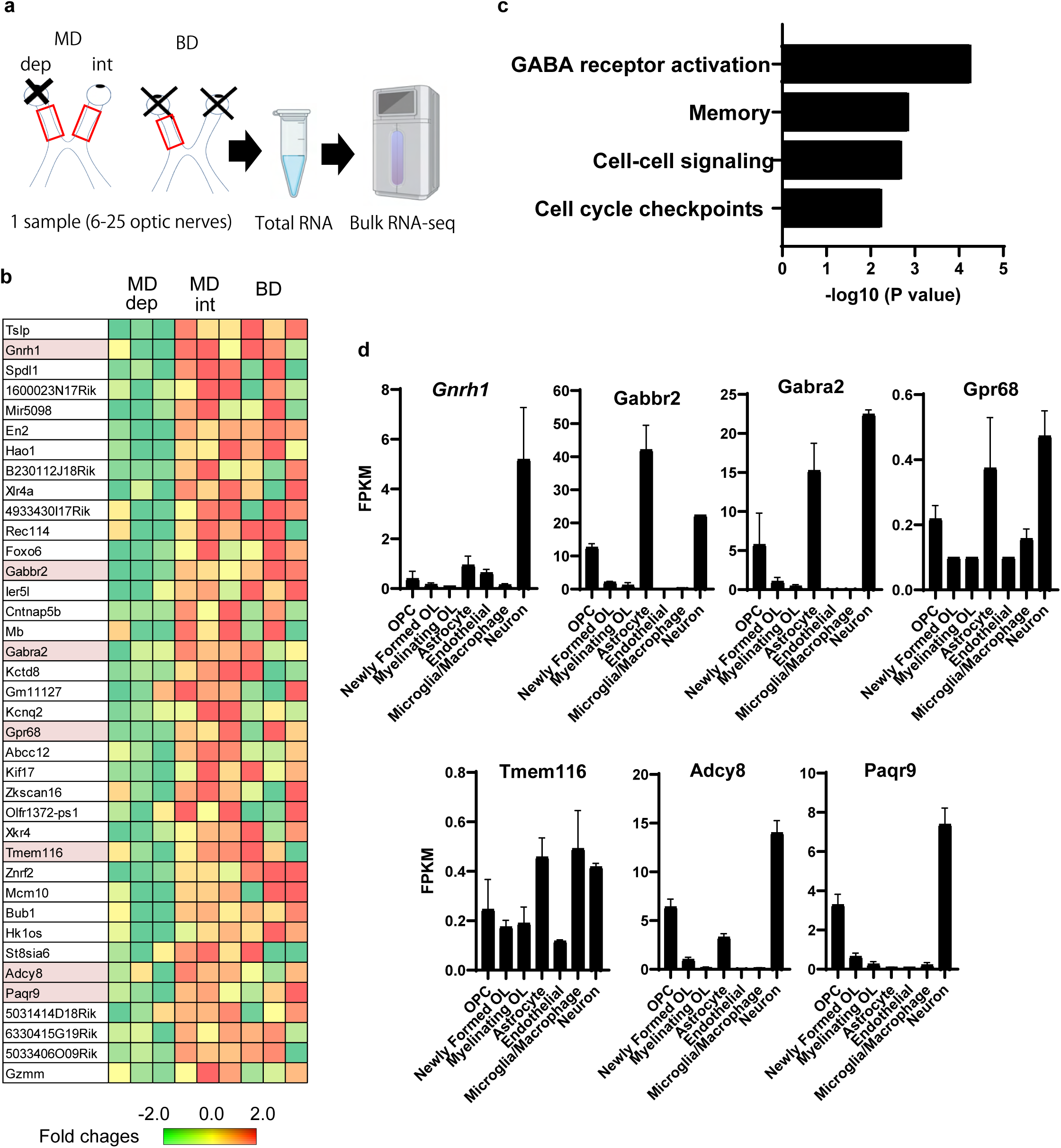
RNA-seq revealed that GABAergic signaling is down regulated in MD deprived side. **a**, Experimental design (drawing created using BioRender). **b**, A heatmap and mean fold change (log_2_ fold change) of genes down regulated in MD deprived optic nerves compared to MD intact and BD optic nerves. Genes related to GABA receptor activation are highlighted by pink. **c**, A result of Gene oncology analysis for genes down regulated in MD deprived optic nerve performed using Metascape website (https://metascape.org). **d**, RNA-seq data of GABA receptor activation genes extracted from Brain RNA-seq database (https://brainrnaseq.org).

Given that GABA receptor activation, cell-cell signaling, and cell cycle check points related genes are down regulated, we hypothesized that oligodendrocytes receive GABAergic inputs that regulate differentiation of OPCs to oligodendrocytes.

To investigate whether GABAergic input could alter the oligodendrocyte differentiation and myelin sheath length in the visual pathway, we administered GABA-A receptor antagonist bicuculline into *PDGFRa-CreERT2:Tau-lox-STOP-lox-mGFP* mice, which label newly differentiated oligodendrocytes with membrane targeted GFP after tamoxifen administration^32^. Tamoxifen was administrated at P21, bicuculline was administrated from P22 to 28, and mice were sacrificed at P30 (Figure 4**a**). We found that the number of newly generated oligodendrocytes (GFP and CC1 positive) were approximately 10 times higher in bicuculline treated optic nerve (Figure 4**b**). The total CC1 positive oligodendrocytes was also higher in bicuculline-treated mice, further confirming excessive oligodendrocyte generation in the bicuculline-treated mice (Figure 4**c** and 4**d**). Since the number of GFP-labeled oligodendrocytes was low in vehicle treated mice, tamoxifen was administrated at P16 to increase the number of GFP-labeled oligodendrocytes and analyze their morphology. We found that the myelin sheath length of bicuculline-treated mice was significantly shorter compared to vehicle treated mice (Figure 4**d** and 4**e**). The number of Caspr signals was significantly higher in bicuculline treated mice (Figure 4**d** and 4**f**). Taken together, the inhibition of the GABA-A receptor increases the number of oligodendrocytes and decreases myelin sheath length in the optic nerve, mimicking abnormal myelination observed in MD deprived side (Figure 4**g**).

**Figure 4.**
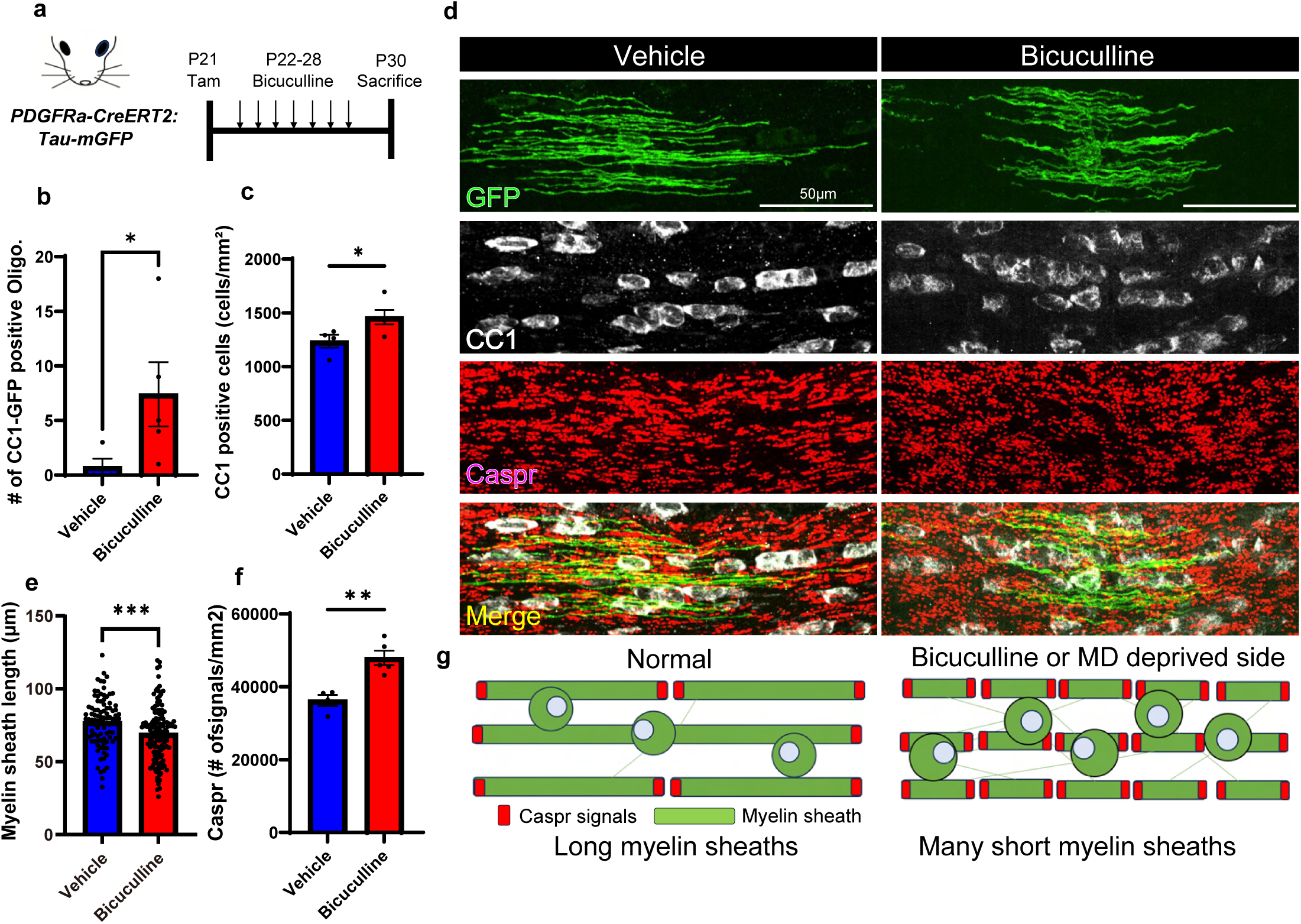
Inhibition of GABAergic signaling mimicks abnormal myelination observed in MD deprived side. **a**, Scheme of the experimental design for administration of bicuculline, GABA-A receptor antagonist, into *PDGFRa-CreERT2:Tau-lox-STOP-lox-mGFP* mice. **b**, The number of GFP-CC1-positive newly generated oligodendrocytes in the optic nerve (within 1 mm from the optic chiasm) in vehicle- and bicuculline-treated mice (vehicle 0.75 ± 0.75 cells, bicuculline 7.4 ± 2.94 cells, optic nerve 1 mm of the optic chiasm, p = 0.032). **c**, The density of CC1-positive mature oligodendrocytes in the optic nerve in vehicle- and bicuculline-treated mice (vehicle 1236 ± 60.2 cells/mm^2^, bicuculline 1459 ± 67.0 cells/mm^2^, p = 0.046). **d**. Representative z-stacked confocal images (20 slices, equivalent to a 6µm thick section) of GFP-positive oligodendrocytes, CC1, and Caspr immunofluorescence in the optic nerves of vehicle- and bicuculline-treated mice. **e**, The myelin sheath length in the optic nerve of vehicle- and bicuculline-treated mice (vehicle 77.3 ± 1.8 µm, bicuculline 69.1 ± 1.6 µm, p = 0.0003). **f**. The density of Caspr signals in the optic nerve of vehicle- and bicuculline-treated mice (vehicle 36225 ± 1532 signals/mm^2^, bicuculline 47938 ± 1975 signals/mm^2^, p = 0.0029). **g**, Schematic summary showing MD and bicuculline treatment produce excessive oligodendrocytes/myelin sheaths and each myelin sheath becomes shorter coinciding with excessive Caspr signals. Data were analyzed by Mann-Whitney U test (**b**, **e**) or by unpaired t test (**c**, **f**). ns *P* > 0.05; \**P* < 0.05; \*\**P* < 0.01; \*\*\**P* < 0.001. Bar graphs show mean values, and error bars indicate SEM.

Next, we investigated whether increasing the GABAergic signaling could ameliorate the excessive oligodendrocytes number and shorter myelin sheaths in MD deprived optic nerve. For this purpose, *PDGFRa-CreERT2:Tau-lox-STOP-lox-mGFP* mice underwent MD and tamoxifen was administrated at P21. Then, tiagabine, GABA reuptake inhibitor that enhances GABAergic signaling, was administrated to these mice from P22 to 28 in accordance with a previous report^33^, and the mice were sacrificed at P30 (Figure 5**a**). The number of newly generated oligodendrocytes double-positive for GFP and CC1 on the MD intact side was similar in vehicle- and tiagabine-treated mice (Figure 5**b** and 5**c**). By contrast, tiagabine administration significantly reduced the number of newly generated oligodendrocytes (Figure 5**b** and 5**d**) and mature oligodendrocytes (Figure 5**e**) in MD deprived side, indicating excessive oligodendrocyte generation on MD deprived side was reduced by tiagabine treatment. In addition, tiagabine administration increased myelin sheath length (Figure 5**f** and 5**g**) and reduced the number of Caspr signals in MD deprived side (Figure 5**h**), compared to vehicle-treated mice, indicating that short and excessive myelin sheaths were rescued by the tiagabine-treatment (Figure 5**f**). Consistently, the difference of myelin sheath length between MD intact and deprived axons was ameliorated by tiagabine treatment, as the ratio of myelin sheath length in MD deprived axons to that of MD intact axon in tiagabine treated mice was significantly higher than that in vehicle-treated mice (Figure 5**i**). Collectively, diminished GABAergic signaling is responsible for myelin abnormality seen in the MD deprived side and that was rescued by enhancing GABAergic signaling.

**Figure 5.**
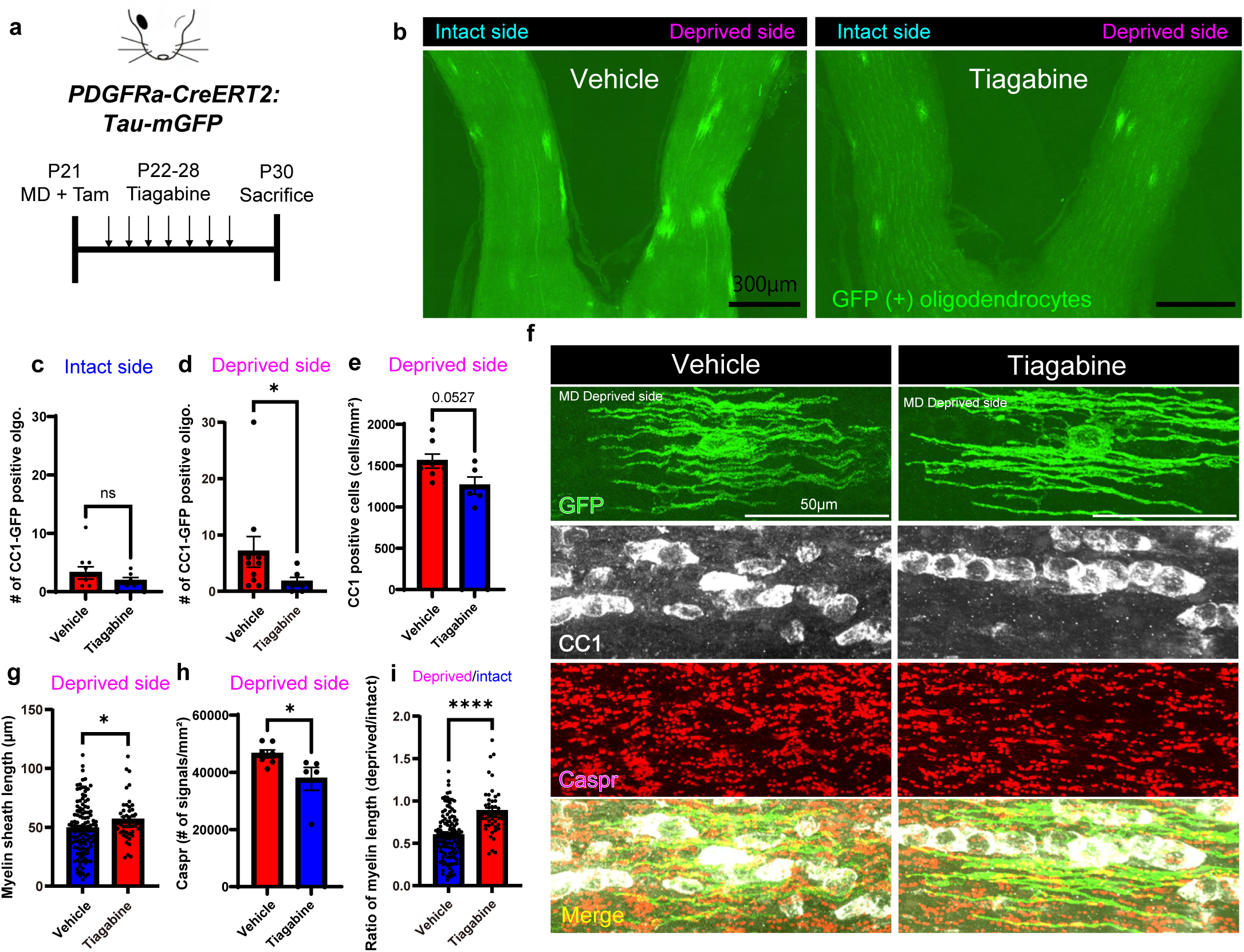
Enhancing GABAergic signaling rescued impaired myelination in the MD mice. **a**, Experimental design for tiagabine administration to enhance GABAergic signaling. **b**, Representative confocal images of MD intact and deprived optic nerves in vehicle- and tiagabine-treated *PDGFRa-CreERT2:Tau-lox-STOP-lox-mGFP* mice subjected to MD. The GFP-positive cells are oligodendrocyte lineage cells differentiated after tamoxifen injection at P21. **c**,**d**, Comparison of the average number of GFP-CC1-positive newly generated mature oligodendrocytes in the MD intact side (**c**) (vehicle 3.2 ± 1.0 cells, tiagabine 1.8 ± 0.6 cells, optic nerve 1 mm rostral to the optic chiasm, p = 0.50) or deprived side (**d**) (vehicle 7.0 ± 2.7 cells, tiagabine 1.7 ± 0.8 cells, optic nerve 1 mm of the optic chiasm, p = 0.034) of vehicle- or tiagabine-treated *PDGFRa-CreERT2:Tau-lox-STOP-lox-mGFP* mice. **e**, Density of CC1-positive cells in MD deprived side of vehicle- or tiagabine-treated mice (vehicle 1555 ± 86 cells/mm^2^, tiagabine 1257 ± 106 cells/mm^2^, p = 0.0527). **f**, Z-stacked confocal images (20 slices, equivalent to a 6 µm thick section) of GFP-CC1-positive cells with Caspr immunofluorescence in the MD deprived optic nerve of vehicle-(left) or tiagabine-treated (right) *PDGFRa-CreERT2:Tau-lox-STOP-lox-mGFP* mice subjected to MD. **g**, Comparison of the myelin sheath length between MD deprived optic nerve of vehicle- and tiagabine-treated *PDGFRa-CreERT2:Tau-lox-STOP-lox-mGFP* mice (vehicle 49.0 ± 2.2 µm, tiagabine 56.3 ± 2.6 µm, p = 0.047). **h**, Density of Caspr signals in MD deprived side of vehicle- and tiagabine-treated mice (vehicle 46434 ± 1387 signals/mm^2^, tiagabine 37717 ± 4031 signals/mm^2^, p = 0.0416). **i**, Comparison of the ratio of myelin sheath length on the MD deprived and intact side, calculated by dividing the individual myelin sheath length of the MD deprived optic nerve by the mean myelin sheath length of the MD intact side in vehicle- or tiagabine-treated *PDGFRa-CreERT2:Tau-lox-STOP-lox-mGFP mice* (vehicle 0.59±0.03, tiagabine 0.88 ±0.05, p < 0.0001). Data were analyzed by Mann-Whitney U test (**c**, **d, g, i**) or by unpaired t test (**e**, **h**). *P* value is indicated in on the graph in (**e**). ns, *P* > 0.05; \**P* < 0.05; \*\*\*\**P* < 0.0001. Bar graphs show mean values, and error bars indicate SEM.

### GABAergic neurons in the optic nerve

To find the source of GABAergic inputs in the optic nerve, whole optic nerve immunostaining for GABA transporter VGAT was performed. We found that VGAT signals formed a belt-like structure in the optic nerve (Figure 6**a**). The number of VGAT belts is approximately one in the 0-1mm range rostrally from chiasm (1.1 ± 0.1 belts, n = 10 optic nerves). The VGAT signals were close proximity to oligodendrocyte lineage cells (Olig2-positive, CC1 weakly positive cells, Figure 6**b** and 6**c**). Furthermore, inhibitory postsynaptic protein gephyrin were expressed in proximity to the presynaptic VGAT signals, suggesting that the oligodendrocyte lineage cells form synapse-like structures and may receive GABAergic inputs (Figure 6**d**). VGAT signals were also present in the vicinity of astrocytes (Olig2-negative, S100B-positive cells) (Figure 6**e**).

**Figure 6.**
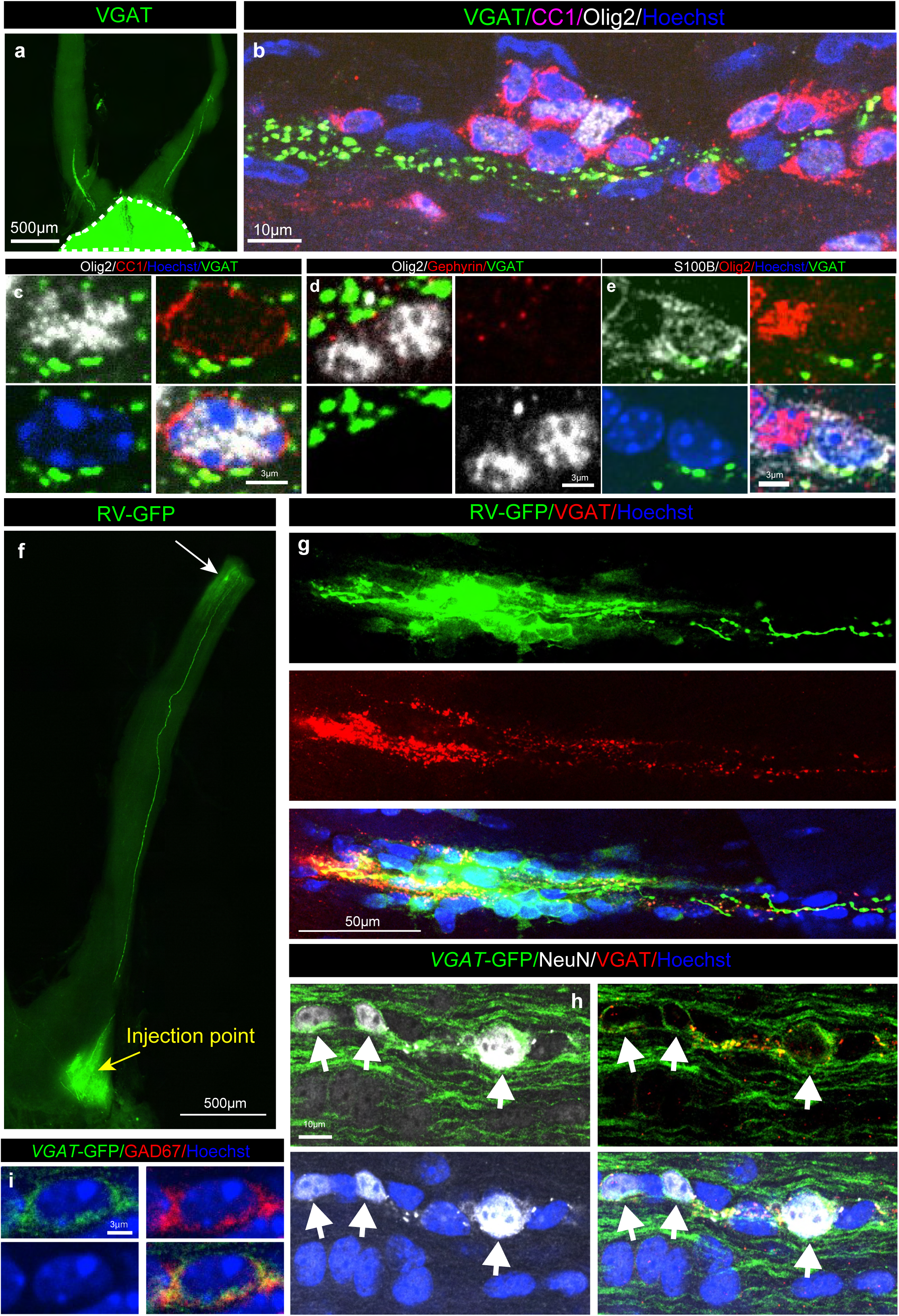
GABAergic neurons in the optic nerve. **a**, VGAT immunofluorescences in the optic nerve. The region surrounded by white broken line indicates the suprachiasmatic nucleus. **b**, VGAT, CC1, and Olig2 immunofluorescences showing VGAT-positive GABAergic terminals were close proximal to CC1-Olig2-positive oligodendrocyte lineage cells. **c**, Representative confocal images of a CC1-Olig2-positive cell surrounded by VGAT signals. **d**, Gephyrin immunofluorescence expressed in Olig2-positive oligodendrocyte lineage cells. **e**, Representative confocal images showing that a S100B-positive-Olig2-negative astrocyte was close proximal to VGAT signals. **f**, A confocal image showing the injection point of retrograde tracer RV-GFP (yellow arrow) and a GFP-positive cell (white arrow) in the optic nerve. **g**, Representative images of RV-GFP positive cell co-localized with VGAT signals. **h**, Representative confocal images of GFP, NeuN, and VGAT signals in the optic nerve of *VGAT-Cre:Tau-lox-STOP-lox-mGFP* mice. **i**, Representative confocal images of GAD67 and GFP signals in the optic nerve of *VGAT-Cre:Tau-lox-STOP-lox-mGFP* mice, showing the presence of a GABAergic cell in the optic nerve.

To identify the origin of GABAergic inputs, we focused on RGC neurons projecting to the suprachiasmatic nucleus, as these neurons are known to include inhibitory RGC neurons^34^. A retrograde neural tracer, attenuated rabies virus encoding GFP^35,36^, was injected into the suprachiasmatic nucleus (Figure 6**f**). Unexpectedly, we found that GFP positive cell bodies were present within the optic nerve (Figure 6**f**), and their neurite processes are well merged with the VGAT signals (Figure 5**g**). To confirm whether these cells are VGAT-positive GABAergic neurons, we used *VGAT-Cre:Tau-lox-STOP-lox-mGFP* mice, where label VGAT expressing cells with the membrane targeted GFP. In the optic nerve, we found that GFP-NeuN double positive cells were present proximal to the VGAT signals (Figure 6**h**). The VGAT-positive cells also expressed GAD67 (Figure 6**i**), indicating that these optic nerve neurons produce GABA. These results indicate that oligodendrocyte lineage cells receive GABAergic inputs from the neurons located in the optic nerve.

## Discussion

It has been shown that neuronal activity directly affects myelination. However, we found that impairment of myelination in optic nerve is not correlated to degree of visual input reduction, MD reduces myelination, but BD does not. We found that abnormal myelination in MD deprived side is caused by reduced GABAergic signaling. Furthermore, we identified GABAergic neurons in the optic nerve as a potential source of GABAergic inputs. Although white matter GABAergic neurons are known to exist in the white matter of human neocortex and called “interstitial neurons” ^37^, to the best of our knowledge, this is the first report that identifies the neurons in the optic nerve. Our results indicate that myelination of the optic nerve is regulated by the balance of visual inputs from both eyes, similar to neuronal plasticity in the visual cortex, via GABAergic signaling.

It remains unclear how cells in the optic nerve know the difference of visual inputs between the eyes. Suprachiasmatic nucleus receive projection from both eyes, and reciprocal synapses are present^38^. It may be possible that optic nerve GABAergic neurons, which project to the suprachiasmatic nucleus, receive inputs from the suprachiasmatic nucleus. One possible mechanism underlying experience-dependent myelination in the optic nerve is that visual inputs from both eyes are integrated and computed in the suprachiasmatic nucleus. Subsequently, GABAergic neurons in the optic nerve receive this information from the suprachiasmatic nucleus and regulate GABA expression. As a result, oligodendrocyte differentiation and its morphology are controlled by inputs from both eyes. Further studies are required to reveal characteristics of GABAergic neurons in the optic nerve and to understand the mechanisms that detect the differences in visual inputs between two eyes and modulate myelination in the optic nerve.

Etxeberria et al (2016) reported that reducing glutamatergic signals shortens myelin sheath length in the optic nerve^13^. In this study, we have shown that enhancing GABAergic signals rescued shorter myelin sheaths in MD mice. Thus, the balance of excitatory and inhibitory signals may be important for proper myelination. We have shown that artificially enhancing unilateral RGC activity by DREADD during BD, which mainly enhances excitatory RGC, lengthens the myelin sheath, but fails to recapitulate shorter myelin sheath on the less active side as seen in MD mice. This result supports the concept that the myelination of the optic nerve is regulated by a fine balance between excitatory and inhibitory signals.

Interestingly, changes in myelination in the optic nerve, ocular dominance in the primary visual cortex, and visual acuity are well correlated. Blais et al (2008) reported that aberrant ocular dominance induced by MD can be recovered by subsequent BD in the mouse model, while dark-rearing did not^39^. In this study, we have shown that BD does not affect myelination, while our previous study reported that dark rearing shortens myelin sheath length^14^. Considering the findings of Blais et al. in conjunction with our study, BD preserves vision, whereas dark rearing impairs vison. In addition, perturbed myelination in dark-reared mice and intact myelination in BD mice suggest that light stimulus is necessary for proper myelination.

Treatments for anisometropic amblyopia include patching and atropine eye drop, which temporarily blur vision, encouraging use of amblyopic eye. It is likely that myelination is impaired in patients with anisometropic amblyopia, and that shortened myelin sheaths are cured by patching or atropine eye drops. Future research into GABA-mediated intervention in the myelination of the optic nerve may lead to new treatment strategies for amblyopia.

Axons in the optic nerve are almost fully myelinated, which is different from the axons in the corpus callosum or the cerebral cortex, where more than half of the axons are unmyelinated^40^. To increase the myelin internodal length, it is necessary to reduce the number of myelinating oligodendrocytes or its producing myelin sheath number. Our data indicate that GABAergic signaling inhibits oligodendrocyte generation in the optic nerve. In MD deprived side, mature oligodendrocyte generation was extended and oligodendrocytes produced shorter myelin sheaths. The shorter internodal length decreases conduction velocity^41^. It is known that MD alters neuronal responses in visual cortex^16,42^. Although the visual acuity of MD deprived eye is significantly reduced^21^, visual functions such as tracking objects increase in MD intact eye^43^. It is possible that the short myelin on the MD deprived side may be important for preferential processing of visual information from the intact eye.

It has been recognized that optic nerve consists of RGC axons and non-neuronal cells^44^. In this report, we have shown that there are GABAergic neurons in the optic nerve. This cell seems to be specialized to modulate oligodendrocyte differentiation and its morphology. This neuron is characterized by its many presynaptic terminals. The neuro-glia forming cells (NGF cells) are GABAergic neurons that are present in the brain and possess many presynaptic terminals^45,46^. It is possible that the optic nerve GABAergic neurons share similar characteristics with NGF cells. NGF cells release GABA into the extracellular matrix from the densely spaced presynaptic terminals, some without postsynaptic specification, which allow GABA to reach both synaptic and non-synaptic receptors located at a distance from release sites as a volume transmission. It is possible that optic nerve GABAergic neurons also release GABA into the extracellular space and activate GABA receptors in this manner.

In conclusion, we revealed that GABAergic signaling modulates oligodendrocyte number and its morphology, likely through optic nerve GABAergic neurons. Investigation of GABAergic signaling-mediated myelination and GABAergic neurons in the other white matter, such as the corpus callosum, will be important to confirm whether this GABAergic regulation of oligodendrocytes is generally essential for activity-dependent myelination. Further studies are required for revealing the mechanism and functional consequence of the activity-dependent myelination through GABAergic signaling.

## Materials and Methods

### Animals

All animals were housed in a cage with 12 h light/dark cycle, and one cage contains one to six adult mice or a mother and one to seven pre-weaned mice. All animal care and procedures were conducted in accordance with guidelines approved by the Ethics Review Board of the Jichi Medical University. All mice were C57BL/6 background, and both female and male mice were used.

*PDGFRa-CreERT* mice^47^ gifted by William D Richardson (University College London) or *VGAT-Cre* mice^48^ (RBRC10606) obtained from RikenBRC were mated with *Tau-lox-STOP-lox-mGFP* mice obtained from the Jackson Laboratory (#021162)^49^ to generate *PDGFRa-CreERT* (heterozygous)*:Tau-lox-STOP-lox-mGFP* (heterozygous) mice or *VGAT-Cre* (heterozygous)*:Tau-lox-STOP-lox-mGFP* (heterozygous) mice. Intraperitoneal administration of Tamoxifen (10 mg/mL, corn oil) was given to *PDGFRa-CreERT:Tau-lox-STOP-lox-mGFP* mice at P16 or P21 at a dose of 100 mg/kg.

### Monocular and Binocular deprivation

Eyelid suturing was performed in accordance with a previous report^50^. Briefly, margin of mouse eyelids was trimmed by Vannas Scissors (501839, WPI, Florida, USA) and eyelids were sutured by surgical suture with needle (CCR760B2NT, Natsume, Tokyo, Japan) under anesthesia with 2–3% isoflurane. After the suturing, eye ointment containing anti-inflammatory agent (Betamethasone sodium phosphate) and an antibiotic (fradiomycin sulfate), rinderon-A ointment (Shionogi Pharma, Osaka, Japan) was applied on the sutured eye.

### Viral injection

For attenuate rabies virus encoding GFP (RV-GFP) injection into the optic chiasm, mice were anesthetized with intraperitoneal injection of a cocktail of medetomidine (0.3 mg/kg), midazolam (4.0 mg/kg) and butorphanol (5.0 mg/kg) or 2–3 % isoflurane. Mice were placed in a stereotaxic frame (Narishige) with a mouse adaptor. After drilling a hole into the skull, RV-GFP (1 μL, 3.3 x 10^4^ IU, 0.2 mm posterior and 0.0 mm lateral to the bregma, depth of 5.0 – 5.3 mm from pia matter) was injected into the optic chiasm. For injection into the suprachiasmatic nucleus, RV-GFP (330 nL; 3.3 x 10^4^ IU) was injected into the suprachiasmatic nucleus using the same method described above.

For intravitreal AAV injection, the mice were anesthetized with 2–3% isoflurane, and 0.5 or 1.0 µL of AAV solution (1.0 × 10^9^ vg) was injected into the vitreous cavity through a Hamilton syringe with a 33-gauge needle (7634–01 and 7803–05, point 4, 0.375 inches, Hamilton) in accordance with a previous report^51^. The needle was lodged for 30 s before withdrawing for preventing leakage in accordance with previous our report^14^. AAV2-hrGFP, AAV7m8-hrGFP, AAV2-BFP^25^, AAV2-hM3Dq-mCherry (50474-AAV2, addgene), and AAV1-Syn-Cox4-dApex2^52^ were used for this research.

### Drug administration

*PDGFRa-CreERT:Tau-lox-STOP-loxmGFP* mice received daily intraperitoneal injections from P22 to 28 of either 1 mg/kg bicuculline (Selleck) in saline containing 2.7% DMSO or 50 mg/kg tiagabine in saline containing 2.7% DMSO, following a previous report ^33^. Vehicle-treated mice were injected with saline containing 2.7% DMSO. The injection volume was 10 µL per gram of body weight, and all solutions were warmed to 37℃ prior to injection. The weight of tiagabine-treated mice was approximately 10 % lower compared to vehicle-treated mice after 6 consecutive days tiagabine-treatment, (Figure S3) as indicated in FDA drug databases as a side effect^53^. Bicuculline-treated had no effect on body weight.

Mice injected with AAV2-hM3Dq-mCherry at P10 were intraperitoneally administrated 10 µL per gram of body weight of clozapine-N-oxide (0.5mg/ml, diluted in saline, 4936/10, Funakoshi) from P24 to 33 (10 consecutive days).

### DAB staining, electron microscopic preparation, and TEM observation

Mice injected with AAV1-*Syn-Cox4-dApex2* were perfused transcardially with a mixture of 2.5% glutaraldehyde and 2% paraformaldehyde in 0.1 M phosphate buffer (pH 7.4) under deep anesthesia. The tissue composed of the optic nerve and optic chiasm were collected and incubated in the same fixative for 2 h at 4℃. The specimens were separated into the optic nerve and optic chiasm, and washed for 10 min with PBS containing 50 mM glycine two times, then washed for 10 min with PBS one time on ice. The specimens were then incubated in the DAB solution (0.3mg/mL in PBS) in the dark with shaking for 2 h RT. Then, 0.3% H_2_O_2_ was added into the DAB solution for final concentration 0.003%. The specimens were incubated in the DAB solution with 0.003% H_2_O_2_ for 8 h in the dark at RT with shaking. Then the specimens were washed for 10 min with PBS, 10 min with PBS containing 50mM glycine, and 10 min with PBS. Then the reduced osmium-thiocarbohydrazide-osmium method (rOTO) staining was performed as described previously^14^. The specimens were then embedded in Quetol 812 epoxy resin (Nisshin EM, Tokyo, Japan) and the resin was incubated at 60℃ to polymerization. Ultrathin sections were prepared with an ultramicrotome (Ultracut UCT, Leica). Images were obtained by TEM (HT7700, Hitachi High-Tech, Tokyo, Japan). The analysis for percentages of myelinated axons, axonal diameter, and G-ratio was performed by blind raters.

### Whole-Mount immunostaining

Whole-mount immunostaining was performed as described in our previous study^14^. Briefly, mice were perfused transcardially with 4% PFA. The eyes or visual pathway containing the optic nerve and optic chiasm were collected and post-fixed with 4% PFA for 15 min at RT or 1 day at 4℃, respectively. The retinae were collected from the eyes after the post-fixation. The fixed specimens were incubated in 2% PBST (2% triton in PBS) for 24-48 h at RT, and the specimens were incubated in the blocking buffer containing 10% normal goat serum at 4℃ for 1–2 days. Then, the specimens were incubated in the following antibodies chicken anti-mCherry (1/500, ab205402, abcam), rabbit anti-c-Fos (1/1000, ab190289, Abcam), rabbit anti-Caspr (1/500, ab34151, Abcam), guinea pig anti-VGAT (1/500, 131004, Synaptic Systems), chicken anti-Olig2 (1/500, OLIG2-0020, Aves), mouse anti-APC (CC1, 1/100, OP80, Merk), rat anti-GFP (1/1000, 04404-26, Nacalai), mouse anti-S100B (1/100, ab34686, Abcam), rat anti-PDGFRa (1/100, CloneAP A5, 558774, BD Biosciences) rabbit anti-Iba1 (1/400, 019-19741, Wako), mouse anti-NeuN (1/100, MAB377, Millipore), and mouse anti-GAD67 (1/100, MAB5406, Millipore) antibodies in an Ab dilution buffer (1% normal goat serum, 0.2% TritonX, 2.5% DMSO, and 0.1% sodium azide in PBS) for 3–4 days at 4℃. After washing with the washing buffer (3% NaCl and 0.2% TritonX in PBS), the specimens were then incubated with goat Alexa Fluor secondary antibodies (1/500, Thermo Fisher) for 2 days at 4℃. After washing, the specimens were incubated with Hoechst (1/2000, 33342, Invitrogen) for 3 h at RT. All incubation steps were conducted on low-speed shakers. After washing with PBS for three times for 1 h each, the specimens were placed on a slide glass and incubated with approximately 35 µL of warmed RapiClear solution for 20 min. Then, the specimens were mounted with RapiClear solution, and the edges of the coverslips were sealed using a nail polish.

### Extraction of total RNA and bulk RNA-seq for optic nerve

Optic nerves were collected from the MD or BD mice under a stereomicroscope immediately after cervical dislocation. Six to 48 optic nerves were placed in 350 µL of RLT buffer (Qiagen) and homogenized by BioMasher II (893062, Nippi, Tokyo, Japan) and passing the samples 7 times through a 26G needle fitted to a disposable syringe. The lysate was added to 300 µL of RLT buffer and inverted and mixed. The lysate was centrifuged at 10,000 x g for 3 min, the supernatant was collected, placed in a gDNA Eliminator spin column and centrifuged at 10,000 x g for 30 s. The flow-through was collected and total RNA was extracted using RNeasy Mini kit as described in our previous study^54^. The total RNA was sent to NGS core at Osaka University for bulk RNA seq analysis.

### Visual cliff test

The closed eyelids of MD or BD mice were opened using Vannas Scissors and all whiskers were trimmed under 2–3 % isoflurane anesthesia with stereotaxic microscopy at P35, more than 3 hr prior to the test. The mice were habituated in the experiment room for over 1 hr. The visual cliff test was conducted in accordance with previous reports with minor modifications^27,28^. The visual cliff apparatus consisted of a wooden box with a rectangular arena (51 × 36 cm^2^) on its upper surface, created using 5-mm-thick Plexiglass and encoded by a white wooden panel to prevent the mouse escape. The arena floor was divided into two areas. In the shallow area (26.5 × 36 cm^2^), checkerboard (2.8 × 2.8 cm white and red squares) is attached directly under the plexiglass. In the second area, deep area (24.5 × 36 cm^2^), the checkerboard was placed at the bottom of the wooden box, 29 cm below from the Plexiglass. The Plexiglass was supported on 1 × 36 cm^2^ props at the shallow and deep ends, hence the actual area is shallow area (25.5 × 36 cm^2^), deep area (23.5 × 36 cm^2^). The illuminance in both areas was set to 850-900 lux to ensure equal brightness. Testing was performed under white noise (Sleepme, marpac, USA), and recorded by video camera (GZ-F270, JVCKENWOOD, Japan).

Each mouse was placed in the shallow area with its head facing away to the shallow/deep border. Behavior was recorded for 5 min after the mouse’s head crossed the border. Time spent in the shallow or deep area was manually measured. Time spent above the prop was excluded from the analysis. A discrimination index was calculated as follows (time spent in shallow area - time spent in deep area)/ total time. The expected value of the discrimination index was 0.04. The number of times the mouse’s head crossed the border was defined as a transition and counted. A retraction was defined and counted when mouse returned to the shallow area immediately after crossing the border with the body parts remaining in the shallow^55^. After each test, the glass and panel were piped first with water and then with 70% ethanol.

### Microscopic analysis

Series of Z-stack projections were acquired at 0.3 or 0.5 µm intervals (objective lens x 60) using Dragonfly high speed confocal microscope system (Oxford instruments, Abingdon-on-Thames, UK) or FV1000 (objective lens x 60) (Olympus, Tokyo, Japan). Images of a figure (Figure S1**c**) was processed with Fiji smooth function for presentation purpose.

### Measurement of internodal length

Length of myelin sheaths in RV-GFP-labelled oligodendrocytes was measured using Z series confocal microscopic images in accordance with our previous study with minor modifications^14^. Myelin sheaths were identified by its running parallel to the axons and the edge of the internode, paranode, that was determined by its round protrusions. The length was measured by Fiji^56^ segmented line function through tracing GFP signals of myelin sheaths from paranode to paranode in the z-series images. The z-series images obtained in our previous study Osanai et al. (2018)^24^ were reanalyzed for internodal length of MD mice. The oligodendrocyte images of BD mice were newly taken for this study. Length of myelin sheaths of GFP-labelled RGC axons with Caspr signals were analyzed as described in our previous study with minor modification. We identified nodes of Ranvier that were marked by a couple of Caspr signals and measured the distance between two adjacent nodes of Ranvier using the Fiji SNT plugin^57^. The measurement of length of myelin sheaths of GFP-labelled RGC axons was performed by blind raters.

Measurement of myelin sheath length in *PDGFRa-Cre:Tau-lox-STOP-lox-mGFP* mice was performed as described in previous studies^13^. Briefly, both edges of myelin internode (paranodes) were identified by Caspr signals and end of GFP-labelled myelin sheaths, and the myelin internodal length was measured by Fiji SNT plugin^57^ by blind raters.

### Statistical analysis

All statistical analyses were performed with Prism 10 (GraphPad Software). The one-way ANOVA with pot hoc Tukey multiple comparisons was used for parametric multiple comparisons. The Kruskal-Wallis test with post hoc Dunn’s test was used for non-parametric multiple comparison. The Mann-Whitney U test or t-test was used for comparing two groups for non-parametric or parametric data, respectively. The boxes and bars in the graphs indicate the mean ± SEM.

## Acknowledgement

We thank Prof. William D. Richardson (University College London) for providing the *PDGFR⍺-CreERT2* mice. We thank Prof. Takeo Horie (Osaka University), Dr. Ayumu Inutsuka (Jichi Medical University), and Dr. Fumihiro Niwa (Jichi Medical University) for helpful discussion about experimental designs. We thank Dr. Tom Kouki, Ms. Megumi Yatabe and Ms. Sasikarn Looprasertkul (Jichi Medical University) for their technical assistance. This study was supported by KAKENHI Grants from the Japan Society for the Promotion of Science, (21H05241, 21H04786, 24H00583 and 20KK0170) to NO, (21K15197, 24K09670, 22H04922 (AdAMS)) to YO, and a research grant from the National Center of Neurology and Psychiatry (No. 3–5) to NO., a research grant from The Uehara Memorial Foundation (202210148) to YO.

---

**Supplemental Figure1.**
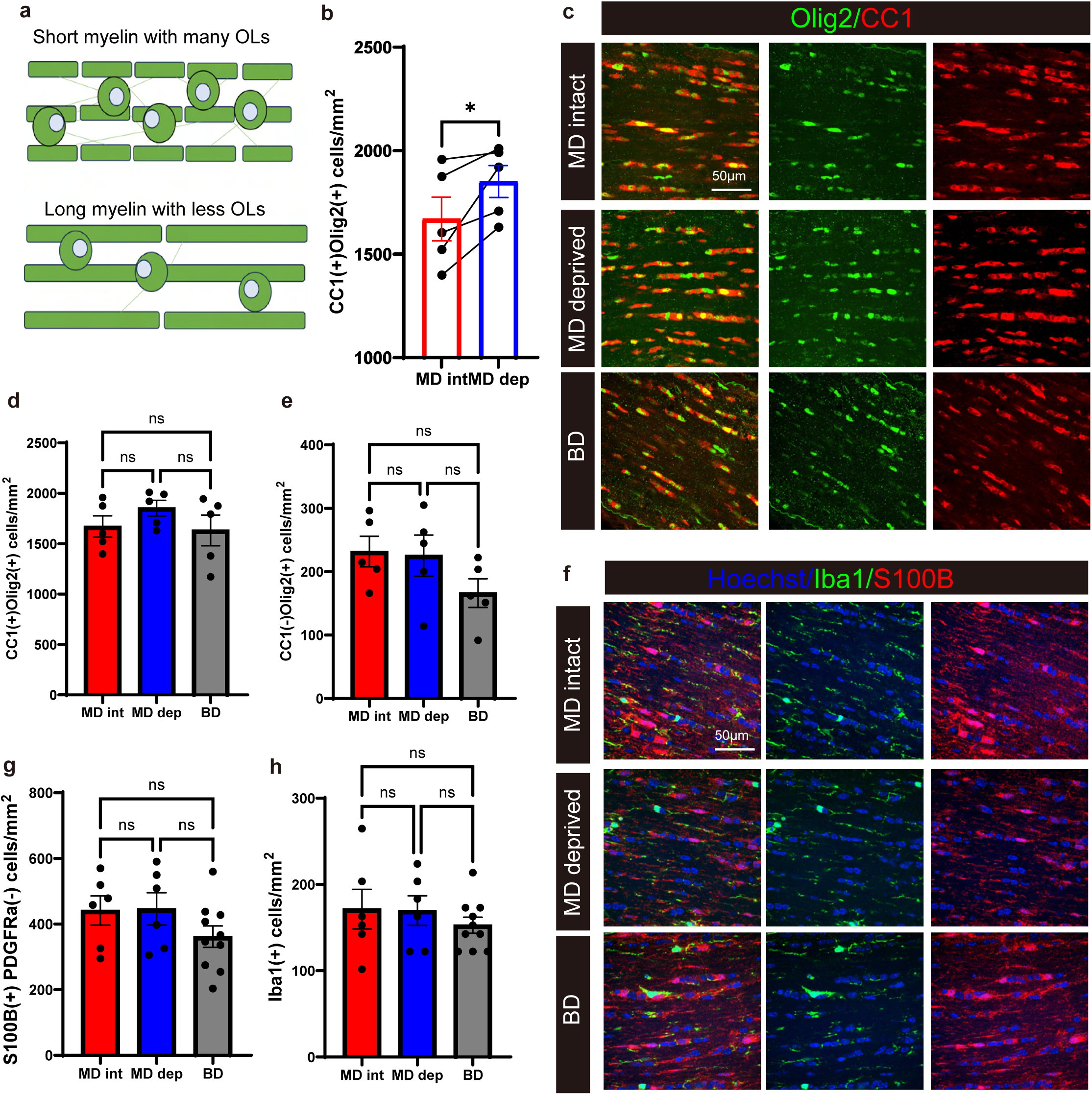
The number of oligodendrocytes in the optic nerve of BD mice is similar to intact optic nerve of MD mice. **a,** Schematic representation of myelination in the optic nerve. Since the optic nerve is fully myelinated, a higher number of oligodendrocytes results in shorter myelin sheath lengths per cell, and vice versa. **b**, Comparison of CC1-Olig2 double positive cell density in the optic nerve between MD intact and deprived sides (MD intact, 1671 ± 106.0 cells/mm^2^, MD deprived 1851 ± 77.2 cells/mm^2^, p = 0.049). **c**, Representative confocal images of CC1- and/or Olig2-positive cells in the optic nerve of MD intact side, MD deprived side, and BD. **d**, Density of CC1-Olig2-double positive cells in the optic nerve of MD intact side, MD deprived side, and BD (MD intact, 1671 ± 106.0 cells/mm^2^, MD deprived 1851 ± 77.2 cells/mm^2^, BD 1632 ± 151.6 cells/mm^2^, p = 0.389). The values of MD intact and deprived side are identical to that of (**b**). **e**, Density of CC1-negative/Olig2-positive cells, in the optic nerve of MD intact side, MD deprived side, and BD (MD intact, 231.8 ± 24.1 cells/mm^2^, MD deprived 225.3 ± 32.5 cells/mm^2^, BD 166.2 ± 22.7 cells/mm^2^, p = 0.204). **f**, Representative confocal images of Iba1- and S100B-positive cells in the optic nerve of MD intact side, MD deprived side, and BD. **g**, Density of S100B-positive/PDGFRa-negative astrocytes (MD intact, 441.1 ± 44.4 cells/mm^2^, MD deprived 446.2 ± 49.1 cells/mm^2^, BD 361.4 ± 32.7 cells/mm^2^, p = 0.239). **h**, Density of Iba1-positive microglia (MD intact, 171.4 ± 23.0 cells/mm^2^, MD deprived 169.7 ± 17.2 cells/mm^2^, BD 152.7 ± 9.2 cells/mm^2^, p = 0.608). Data were analyzed by paired t test (**b**) or by one-way ANOVA with Tukey post hoc test (**d**, **e**, **g**, **h**). ns *P* > 0.05; \**P* < 0.05. Bar graphs show mean values, and error bars indicate SEM.

**Supplemental Figure2.**
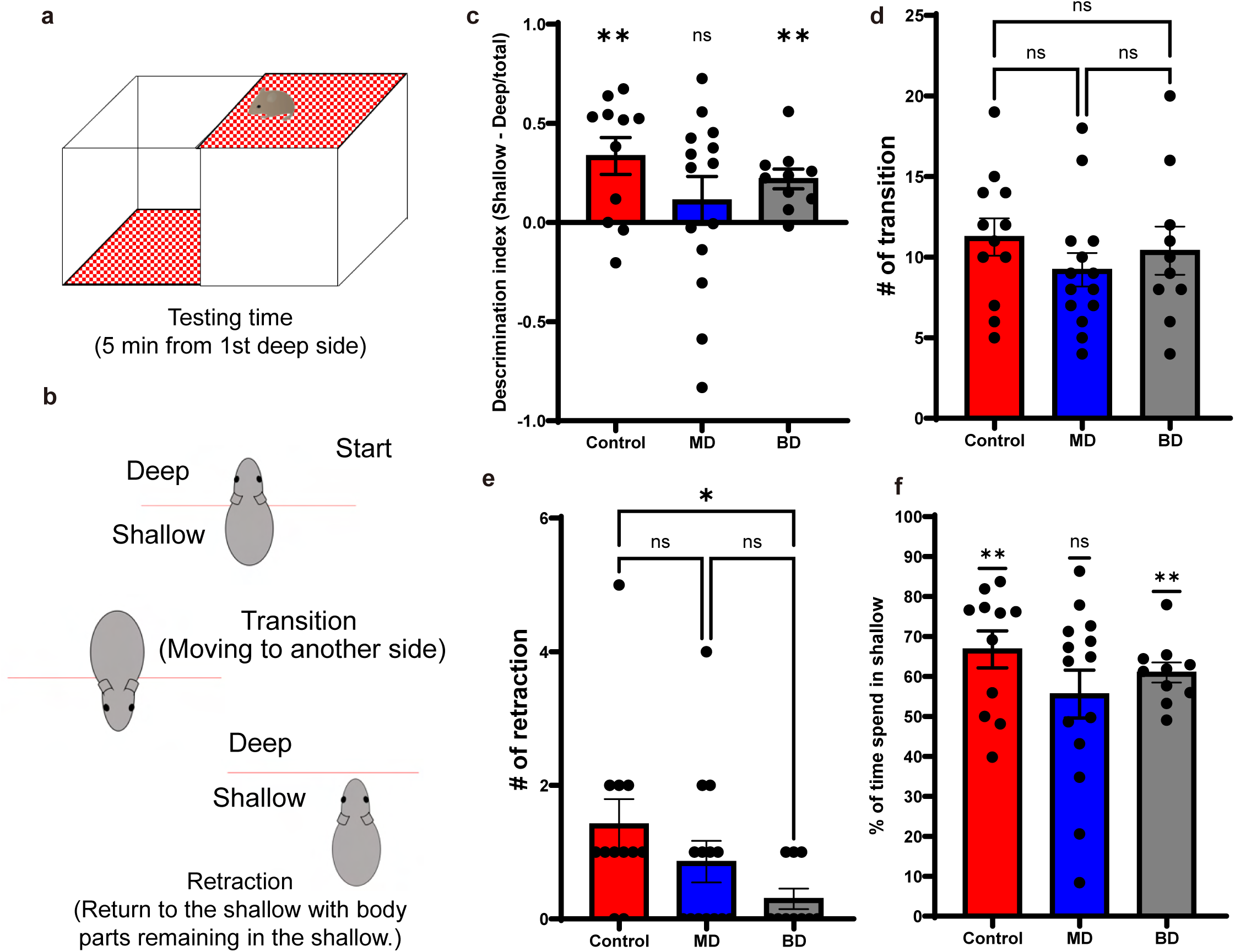
Visual cliff test revealed that depth perception of BD mice is preserved. **a,** Schematic drawing of the visual cliff test. Mice were placed in the shallow area, and their behavior was analyzed for 5 min from the first encounter with the border. **b**, Definition of mouse behaviors. Start: measurements started immediately after the mouse’s head crossed the shallow and deep border. Transition: the transition was counted when the mouse’s head crossed the border. Retraction: the retraction was defined when mice were encountered the border and returned to shallow with body parts remaining in the shallow area. **c**, Comparison of discrimination index between control, MD, and BD mice (discrimination index = (shallow time – deep time)/total time, control 0.34 ± 0.09, MD 0.11 ± 0.12, BD 0.22 ± 0.05, one sample t-test, control p = 0.0098, MD p = 0.56, BD p = 0.0058). **d**, Comparison of the number of transitions between control, MD, and BD mice (control 11.3 ± 1.2, MD 9.2 ± 1.0, BD 10.4 ± 1.5, p = 0.345). **e**, Comparison of the number of retractions between control, MD, and BD mice (control 1.42 ± 0.38, MD 0.86 ± 0.31, BD 0.30 ± 0.15, p = 0.0293). **f**, Comparison of the percentage of time spent in the shallow area (control 66.8 ± 4.6 %, MD 55.6 ± 6.0 %, BD 61.0 ± 6.0 %, control p = 0.0098, MD p = 0.56, BD p = 0.0058). Data were analyzed by one-sample t test (**c, f**) or by Kruskal-Wallis test with post hoc Dunn’s test (**d**, **e**). Expected values for (**c**) and (**f**) are 0.04 and 52%, according to the area of shallow and deep. ns, *P* > 0.05; \**P* < 0.05; \*\**P* < 0.01. Bar graphs show mean values, and error bars indicate SEM.

**Supplemental Figure3.**
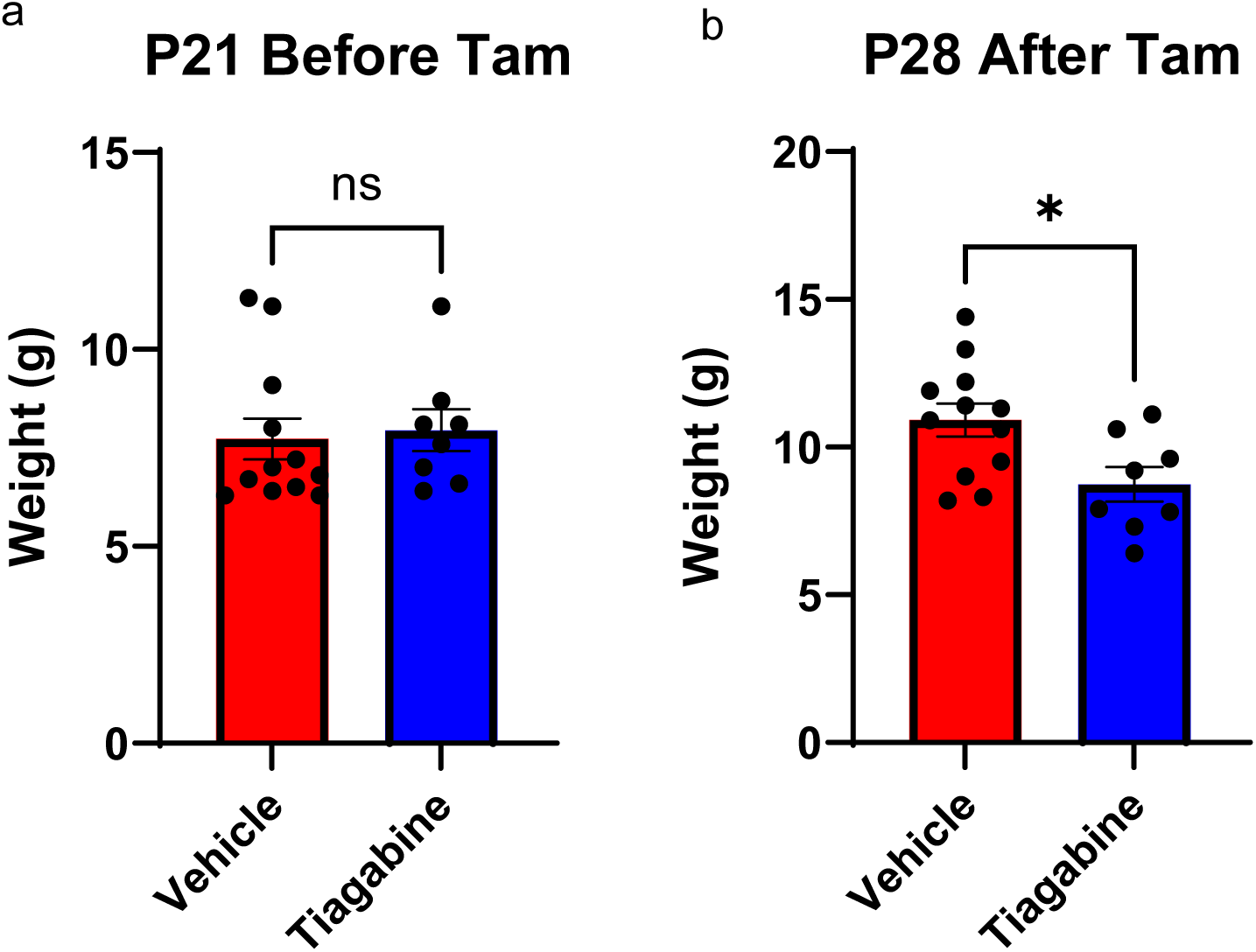
Tiagabine treatment reduces mouse body weight. **a**, Body weight of P21 mice before administration of tiagabine (vehicle 7.73 ± 0.52, tiagabine 7.95 ± 0.53, p = 0.437). **b**, Body weight of P28 mice after 6 consectutive days tiagabine administration or vehicle administration (vehicle 10.92 ± 0.56, tiagabine 8.74 ± 0.58, p = 0.0168). Data were analyzed by unpaired t test. ns *P* > 0.05; \**P* < 0.05. Bar graphs show mean values, and error bars indicate SEM.

